# Acute and Lifelong Exercise Modulate the Tumorigenic Potential of Human Lung Cancer Cells and Their Susceptibility to Cisplatin

**DOI:** 10.64898/2026.03.19.713009

**Authors:** Carlos M. Soares, João P. Moura, Leonardo M. R. Ferreira, Ana Pedrosa, Pedro Filipe, Luís Rama, Ana M. Teixeira, Ana M. Urbano

## Abstract

The association between higher levels of physical activity and lower cancer risk and mortality is well established. However, a causal link is yet to be proven. Recent studies showed a decrease in the proliferation rates of cultured human cancer cells when the human serum employed to stimulate them was conditioned by acute exercise. Here, we tested the hypothesis that serum mediates some of the putative benefits of exercise on cancer through alterations to the growth pattern and susceptibility to chemotherapy agents of cancer cells. To this end, human non-small cell lung cancer (NSCLC) cells were exposed to serum from two cohorts that differed significantly on their levels of physical activity and, accordingly, cardiorespiratory fitness, but were otherwise identical (master athletes and non-exercisers), collected before and after an acute exercise intervention. Serum levels of glucose, lipids, albumin, C-reactive protein and cytokines were determined and the impact of the serum responses to acute and lifelong exercise on the above-mentioned parameters were analyzed. We found that acute exercise decreased the cells’ proliferation rate, yet shortened the cells’ lag phase after detachment, whereas lifelong exercise had the opposite effects. Significantly, we showed, for the first time, that lifelong exercise increased susceptibility to a chemotherapy agent (cisplatin), which may contribute to the decreased cancer mortality rates found among those who exercise regularly. Similar to the cellular effects, changes to serum cytokine levels – several of them linked to the senescence-associated secretory phenotype – depended on whether serum was conditioned by acute or by chronic exercise.

**Key points:**

Chronic exercise increased the in vitro susceptibility of lung cancer cells to cisplatin.
Acute and chronic exercise modulated the in vitro tumorigenic potential of lung cancer cells.
Effects were mediated by serological changes produced by exercise.
Acute and chronic exercise had distinct impacts on serological cytokine levels.

## Introduction

Cancer is one of the world’s largest health problems, accounting for every sixth death globally. And yet, a large percentage of cancer cases and cancer deaths could be prevented simply by eliminating or reducing exposure to environmental and occupational risk factors (Urbano *et al*., 2008; Urbano *et al*., 2012; Katzke *et al*., 2015; Lewandowska *et al*., 2019; Adam *et al*., 2023) and by adopting healthier lifestyles, namely quitting smoking, maintaining a healthy weight, reducing alcohol consumption, and being more active (Mctiernan, 2008; Friedenreich *et al*., 2010; Garcia & Pearce, 2023).

Up to the early 2000s, robust evidence for an association between physical inactivity and increased cancer risk was restricted to a few site-specific cancers (Vainio *et al*., 2002; Friedenreich *et al*., 2010). In the meantime, a wealth of evidence for many other cancer types has been gathered. In particular, a pooled analysis from 2016 of 12 prospective cohort studies involving 1.44 million participants showed that the risk for 13 out of 26 common types of cancer was lower for those participants who self-reported engagement in higher levels of moderate or vigorous intensity leisure-time physical activity, than for those participants who self-reported lower levels of engagement (Moore *et al*., 2016). Specifically, the association was strong (greater than 20% reduction in risk) for oesophageal adenocarcinoma, myeloid leukaemia, and cancers of the liver, lung, kidney, gastric cardia and endometrium, and moderate (10%–20% reduction in risk) for myeloma and cancers of the colon, head and neck, rectum, bladder, and breast(Moore *et al*., 2016).

The evidence for a positive association between exercise and reduced cancer mortality has been very recently strengthened by a randomized trial, conducted at 55 centres and involving a total of 889 patients with resected colon cancer, showing that a 3-year structured exercise program initiated soon after adjuvant chemotherapy significantly extended disease-free survival, with findings consistent with longer overall survival (Courneya *et al*., 2025).

Along the same lines, evidence also emerged for a relationship between the amount of exercise and the reduction in the risk of some cancer types (Thune & Furberg, 2001). A very recent meta-analysis of 96 reported studies involving over 30 million participants yielded an inverse non-linear dose–response association between non-occupational physical activity and disease and mortality outcomes for cancer (total and site-specific), suggesting more pronounced increases in potential benefits from inactive lifestyles up to levels of non-occupational physical activity equivalent to the minimum level recommended by the World Health Organization (WHO) for adults (*i.e.*, “at least 75 to 150 minutes of vigorous-intensity aerobic physical activity per week, or at least 150 to 300 minutes of moderate-intensity aerobic physical activity per week, or an equivalent combination throughout the week” (Bull *et al*., 2020)). Benefits became incrementally smaller up to twice the recommended minimum level, the evidence for benefit being weaker thereafter (Garcia & Pearce, 2023).

Despite the amount and strength of the evidence just discussed, these associations are not proof that higher levels of physical activity reduce cancer risk and improve cancer management. For instance, it cannot be ruled out that the lower cancer risk found amongst active people is a consequence of other healthy lifestyle behaviours besides being physically active or a result of gene variants enriched in active people. Whether a causal relationship exists can only be ascertained if the cellular and molecular mechanisms behind the putative benefits of active lifestyles to cancer prevention and management are firmly established.

It has been argued that the observed reduction of cancer risk and mortality associated with physical activity is primarily an indirect effect of weight loss, especially in terms of adiposity. The fact that an association was found between excess body fat and a higher risk for a large number of cancer types, most notably cancers of digestive organs and cancers of hormone-sensitive organs in women (Lauby-Secretan *et al*., 2016; Avgerinos *et al*., 2019), lends some support to this view. Lowering the number of adipose cells and/or their size reduces the secretion and, hence, the circulating levels of several biomarkers of cancer risk, namely sex hormones, metabolic hormones (*e.g.*, leptin and resistin), insulin-like growth factors, pro-inflammatory cytokines, and macrophage infiltration into adipose tissue, ultimately lowering systemic inflammation (Avgerinos *et al*., 2019; Michailidou *et al*., 2022; Tilg *et al*., 2025). In addition, obesity weakens the antitumor responses of natural killer (NK) cells, thus compromising immunosurveillance and the elimination of cancer cells (Michelet *et al*., 2018). Nonetheless, the fact that the inverse associations between physical activity and cancer risk were unaffected by body size in 10 of the above-mentioned 13 types of common cancers (Moore *et al*., 2016) suggests that mechanisms independent from weight loss contribute to the putative beneficial effects of physical activity to cancer risk and mortality.

There is increasing evidence that the growth patterns of cultured human cancer cells are altered when the human serum used to stimulate them is conditioned by exercise. These *in vitro* studies are still sparse and vary considerably in all aspects of study design, from exercise intervention to cohort characteristics, cell lines used, exposure regimen (percentage of serum in the growth medium and duration of the exposure) and assays employed, making direct comparisons difficult. Nonetheless, meta-analyses suggest that, compared to non-conditioned (baseline) serum, serum conditioned by an acute exercise intervention diminishes the viability of cultured cancer cells, with a large overall effect size that increased with the intensity of the exercise (Orange *et al*., 2020; Brown *et al*., 2021; Soares *et al*., 2021). Of note, no studies on the impact of serological responses to exercise on cancer susceptibility to chemotherapy have been reported. In this study, we exposed A549 human non-small cell lung cancer (NSCLC) cells to human sera obtained before or after acute exercise and analyzed the impact of the transient serum responses to the intervention on the sera’s ability to stimulate cancer cell proliferation, on the cells’ plating efficiency (assessed previously in only one study), here used as a metric for reproductive potential, lag phase (not previously reported), cell migration (not previously reported) and susceptibility to chemotherapy (not previously reported). We used two distinct cohorts with different levels of physical activity, volunteers who did not meet the above-mentioned WHO guidelines on physical activity for health (hereafter termed non-exercisers) and master athletes. Through the use of these two cohorts, we were able to also assess the impact of permanent serum changes on the same parameters, as well as to gain insight into how the impact of acute exercise on serum’s cancer cell modulatory properties depends on levels of physical activity throughout life.

## Materials and Methods

### Study overview and ethics statement

Twenty-six Caucasian healthy male volunteers, aged 40 to 63 years were recruited based on their levels of physical activity and assigned to two age-matched cohorts: 13 non-exercisers (individuals who self-reported not having met in the 20 years that preceded their recruitment the WHO guidelines on physical activity for health (Bull *et al*., 2020)) and 13 age-matched master athletes, all of them self-reported experienced competitors who had been training and competing for at least 20 years in their sports modality at the time of recruitment. Blood was collected from each participant before (baseline serum) and after the exercise intervention (post-exercise serum). Serum prepared from the collected blood was used to evaluate the effects of serum responses to acute and lifelong exercise on the pattern of growth of human lung cancer cells, providing valuable information regarding their transformation degree and, ultimately, their tumorigenic potential, as well as on the susceptibility of these cells to the chemotherapy agent cisplatin. All sera were tested in parallel. For ethical considerations, the amount of blood collected was limited, meaning that pooled sera, instead of individual sera, were used in the assessment of plating efficiencies, susceptibly to cisplatin and cytokine detection. Each of the four pooled sera, corresponding to the four conditions just described, contained equal volumes of sera from 13 participants. At the time of recruitment, the intervention and potential risks were explained to the participants who all gave written informed consent before their inclusion in the study. Prior to the acute exercise intervention, participants were subjected to a brief medical examination that included blood pressure measurement, electrocardiogram and a standard medical history questionnaire. Exclusion criteria included smoking, having donated blood in the three months that preceded the intervention, any evidence of chronic or inflammatory diseases, having had an infectious disease up to six weeks prior to the exercise intervention, and taking supplementation or medication. Participants were asked to maintain their lifestyle routines and to avoid caffeine, exercising and partaking in sports activities in the 24 h preceding the exercise intervention. The study was approved by the Ethics Committee of the Faculty of Sports Sciences and Physical Education of the University of Coimbra (reference CE/FCDEF-UC/00062013). All procedures conformed to the Declaration of Helsinki (World Medical, 2013) and with data protection and security regulations (Harriss & Atkinson, 2015).

### Participant characterization

Height and body weight were determined using, respectively, a Harpenden 98.603 stadiometer (Holtain, Crosswell, UK) and a Seca 770 electronic personal scale (Seca, Hamburg, Germany). Body mass composition was assessed using a InBody 770 tetrapolar bioelectrical impedance analyzer (InBody, Cerritos, California, USA) (Brewer *et al*., 2021). Basal metabolic rate (BMR) was calculated by the analyzer according to Cunnigham equation (BMR (in kcal/day) = 370 + 21.6 × fat-free mass (in kg)) (Cunningham, 1991).

### Acute exercise intervention

All participants were asked to perform a maximal incremental voluntary aerobic protocol (Åstrand, 1964), which was carried out on an electromagnetically braked Excalibur Sport bicycle ergometer (Lode B.V., Groningen, NL) and was preceded by a 5-min warm-up of moderate cycling. Throughout the protocol, expired gases were analysed using a Quark CPET breath-by-breath automated gas-analysis system (COSMED, Rome, Italy), which was also used for continuous telemetric determination of heart rate. Initially set at 75 W, the power output was increased by 25-W increments every 3 min, with participants maintaining a sustained cadence of 85–105 rotations per minute (rpm), until volitional exhaustion was reached (*i.e.*, until participants expressed their inability to continue exercising) or when at least two of the following three criteria were met: (i) no increase in oxygen consumption (VO_2_) despite workload increase; (ii) respiratory exchange ratio (ratio of volume of CO_2_ produced to volume of O_2_ consumed) > 1.10; (iii) heart rate above 90% of the estimated maximal value (Beaver *et al*., 1986; Edvardsen *et al*., 2014). Ratings of perceived exertion using the Borg CR-10 scale (Borg, 1982) were self-assessed at each stage and at the end of the protocol. VO_2_ max, *i.e.*, the maximum rate of O_2_ consumption attained during physical exertion, was considered as the highest of the mean VO_2_ values calculated for the two 30-s periods that preceded the end of the intervention (Beaver *et al*., 1986; Edvardsen *et al*., 2014). Index finger capillary L-lactate levels before and at the end of the exercise intervention were determined using a Lactate Pro2 portable meter (Arkray, Amstelveen, NL).

### Blood collection and serum preparation and storage

At two time points (15 min before the intervention and within 5 min into active recovery), whole blood (10–15 mL) was collected from each participant, immediately transferred to a BD Vacutainer™ SST™ II Advance serum separator tube (Becton Dickinson and Company, Franklin Lakes, NJ, USA) and allowed to clot naturally by leaving it undisturbed, at room temperature. After centrifugation (10 min at 1,600 *g*), the off-the-clot serum was carefully removed and immediately stored at −80 °C, in 0.5 mL aliquots, until use. Before use, sera were thawed and, after homogenization, were filtered through 0.2 µm-pore Millex^®^-GV sterile syringe filters (Merck, Darmstadt, Germany; SLGV033RB).

### Quantification of glucose, cholesterol, high-density lipoprotein cholesterol, low-density lipoprotein cholesterol, triglycerides and albumin

Reagents from BioSystems (Barcelona, Spain) were used for the quantification of glucose (COD 11504), cholesterol (free and esterified; COD 11505), high-density lipoprotein cholesterol (HDL cholesterol; COD 11757), low-density lipoprotein cholesterol (LDL cholesterol; COD 11785), triglycerides (11528) and albumin (COD 11573) in individual sera, according to the manufacturer’s instructions. A human, serum-based general biochemistry calibrator (COD 18044) and human biochemistry control sera (COD 18042 and 18043), also from BioSystems, were used for calibration and to verify the accuracy of the measurements, respectively. All analytes were assayed in duplicate and the two values were averaged.

### Quantification of C-reactive protein

C-reactive protein (CRP) was quantified by turbidimetry using and assay reagent (COD 31321), a calibrator (COD 31113) and a control serum (COD 31213) from Biosystems. All sera were tested in duplicate and the values were then averaged. The procedure was that specified by the manufacturer.

### Culture of A549 cells

All cellular studies were performed in cultures of A549 human lung cancer cells (ATCC CCL-185; RRID:CVCL_0023). These cells were routinely grown at 37 °C in a humidified atmosphere of 5% CO_2_/95% air, in filter-vented flasks (Orange Scientific, Braine-l’Alleud, Belgium; 5520100)) containing *ca.* 0.2 mL/cm^2^ of growth medium (RPMI-1640; Merck; R6504) supplemented with 10% (v/v) heat-inactivated fetal calf serum (FCS; Thermo Fisher Scientific, Waltham, MA, USA; 10270-106).

To investigate the impact of exercise-conditioned human serum on cell proliferation, lag phase and plating efficiency, cells were harvested in serum-free growth medium, centrifuged for 5 min at 200 *g*, to remove traces of FCS, and resuspended in an adequate volume of fresh serum-free medium. A small volume (10 µL) of this serum-free cell suspension was subsequently added to culture plates already containing pre-warmed (37 °C) growth medium supplemented with human serum, to a final serum concentration of 10% (v/v).

To investigate the impact of exercise-conditioned human serum on cell migration and for the determination of half-maximal cytotoxic concentrations (CC_50_ values) for cisplatin, cells were harvested in growth medium supplemented with 10% (v/v) heat-inactivated FCS and new cultures were established in this same medium. At *ca.* 24 h post-seeding, spent growth medium was replaced by growth medium supplemented with 10% (v/v) human serum.

### Assessment of cell proliferation and lag phase

For the assessment of cell proliferation, cultures were prepared in 96-well plates (Orange Scientific; 4430100), at a seeding density of 5 000 cells/cm^2^, in 100 µL of growth medium supplemented with 10% (v/v) human serum. In each independent experiment, a total of six replicate cultures were prepared for each serum tested: three each for the estimation of the number of cells in culture at 24 h and 72 h post-seeding. Cell numbers were estimated using the sulforhodamine B (SRB) assay, essentially as described in (Vichai & Kirtikara, 2006). Briefly, at each of the two time points, cells were fixed using a cold 10% (w/v) trichloroacetic acid (Merck; T6399) solution and were then stained using a 0.05% (w/v) SRB solution (Merck; 230162). After washing four times with a 1% (v/v) acetic acid solution (Merck; A6283), the protein-bound dye was solubilized in 200 µL of a 10 mM Tris base solution, pH 10.5 (Merck; 252859), homogenized and the absorbance of the resulting solutions was read at 550 nm (*A*_550_) against a reagent blank (no cells), using a µQuant microplate reader (BioTek Instruments, Winooski, VT, USA). Proliferation rates were then estimated as the fold increase in *A*_550_ over the 48-h period between the two measurements. *A*_550_ values at 24 h post-seeding was used as a metric for lag phase.

### Assessment of plating efficiency

Plating efficiencies were evaluated by the clonogenic assay (Franken *et al*., 2006). To this end, cells were plated as a single-cell suspension at a colony-forming density of 40 cells per well in 24-well plates (Orange Scientific; 4430300), in 500 µL of growth medium supplemented with 10% (v/v) human serum. After nine days of incubation, each culture was washed twice with 2 mL of phosphate-buffered saline (PBS) and colonies were subsequently fixed and stained for 40 min with 400 μL of an aqueous solution containing 6.0% (v/v) glutaraldehyde (Merck; G5882) and 0.5% (w/v) crystal violet (Merck; 548629). Excess fixing/staining solution was then removed, and cultures were washed several times with water. Finally, wells were photographed, colonies were counted manually, and the area of the wells covered by the colonies was measured by ImageJ software version 1.53e (National Institutes of Health, Bethesda, MD, USA). Colony mean size was calculated by dividing this area by the total number of colonies, and plating efficiency was calculated by dividing the number of colonies formed by the number of cells seeded. Each pooled serum was tested in three replicate cultures.

### Assessment of cell migration

*In vitro* cell migration was estimated employing the wound closure assay (Liang *et al*., 2007; Cappiello *et al*., 2018), using IBIDI’s silicone 3-well culture inserts (IBIDI^®^, Gräfelfing, Germany; 80369) to create two artificial 500 µm cell-free gaps in confluent monolayer cultures, corresponding to two technical replicates. These inserts were placed onto the wells of 24-well plates (Orange Scientific; 4430300), one insert per well and per individual serum tested. Cells were then seeded into the insert wells, at a density of 2.5 × 10^5^ cells/cm^2^, in 70 μL of growth medium supplemented with 10% (v/v) FCS. After 22 h of incubation, the inserts were removed to expose two cell-free gaps in each confluent monolayer, growth medium was aspirated, monolayers were washed twice with PBS and three different regions in each gap were imaged (first time point). Pre-warmed (37 °C) growth medium supplemented with 10% (v/v) human serum (500 µL) was then added, cultures were returned to the incubator and, after 18 h, growth medium was once again aspirated, monolayers were washed twice with PBS and the same regions in the gaps were imaged (second time point). All images were captured with a 40× magnification, using an Olympus CKX53 inverted optical microscope equipped with a camera and the EPview™ software (V2.9.6_20201224; Hachioji, Tokyo, Japan). Cell-free areas in the regions imaged were measured using ImageJ and the percentage of gap closure over the 18-h period was calculated. For each serum tested, the average of the respective six percentage values was used as an estimate of cell migration potential.

### Determination of CC_50_ values for cisplatin

To determine the concentration of cisplatin that causes 50% of cell death (CC_50_) in A549 cultures, cultures were prepared in 96-well plates (Orange Scientific), with a seeding density of 5 000 cells/cm^2^, in 100 µL of growth medium supplemented with 10% (v/v) FCS. After a 24-h incubation, spent growth medium was replaced with pre-warmed (37 °C) growth medium supplemented with 10% (v/v) human serum containing 0–455 μM cisplatin (Merck; 232120) and cultures were further incubated for 72 h. For each pooled serum, the cytotoxicity of each cisplatin concentration was tested in three replicate cultures using the SRB assay (as described above). CC_50_ values were obtained from concentration-response curves generated using a nonlinear regression model in GraphPad Prism 9.0.0. software for Windows (GraphPad Software Inc, Boston, Massachusetts, USA).

### Semi-quantitative detection of cytokines

Semi-quantitative assessment of 80 cytokines was carried out in 0.5 mL samples of pooled sera using the RayBio^®^ C-Series Human Cytokine Antibody Array C5, from RayBiotech (Norcross, GA, USA; AAH-CYT-5-4). Samples of pooled sera were first diluted two-fold with the blocking buffer provided with the kits. Following various blocking, incubation and washing steps, performed according to the manufacturer’s instructions, chemiluminescence was detected using a ChemiDoc MP Imaging System (Bio-Rad, Hercules, CA, USA), after a 30-second exposure. The signal intensity (average pixel/area) for each antigen-specific antibody spot was then determined using ImageJ.

### Statistical analysis

Data were analyzed using GraphPad Prism 9.0.0. software for Windows. Differences between the two cohorts were assessed for statistical significance using Student’s unpaired *t*-test or the Mann-Whitney test, according to the normality of the data, which was verified using the Shapiro-Wilk test. The statistical significance of the differences between the two cohorts in terms of the impact of the acute exercise intervention on serum responses serum-mediated modulation of growth properties and sensitivity to cisplatin cells was assessed using repeated measures two-way ANOVA, assuming sphericity. Whenever the null hypothesis was rejected, Šídák’s multiple comparison test was performed (Glantz *et al*., 2001; Maxwell & Delaney, 2004). Differences were considered significant when *P* < 0.05.

## Results

### Cohort characteristics

Selected anthropometric and physiological characteristics of the two age-matched male cohorts employed in this study are summarized in Table 1. The most pronounced anthropometric differences between the cohorts were related to body fat content, with non-exercisers exhibiting, on average, *ca.* 40% higher fat mass (*P* ≤ 0.001) and 60% higher visceral fat area (*P* = 0.002) than master athletes. Non-exercisers also exhibited *ca.* 10% lower muscle mass (*P* = 0.008) and 5% higher BMI (*P* = 0.026).

**Table 1.**
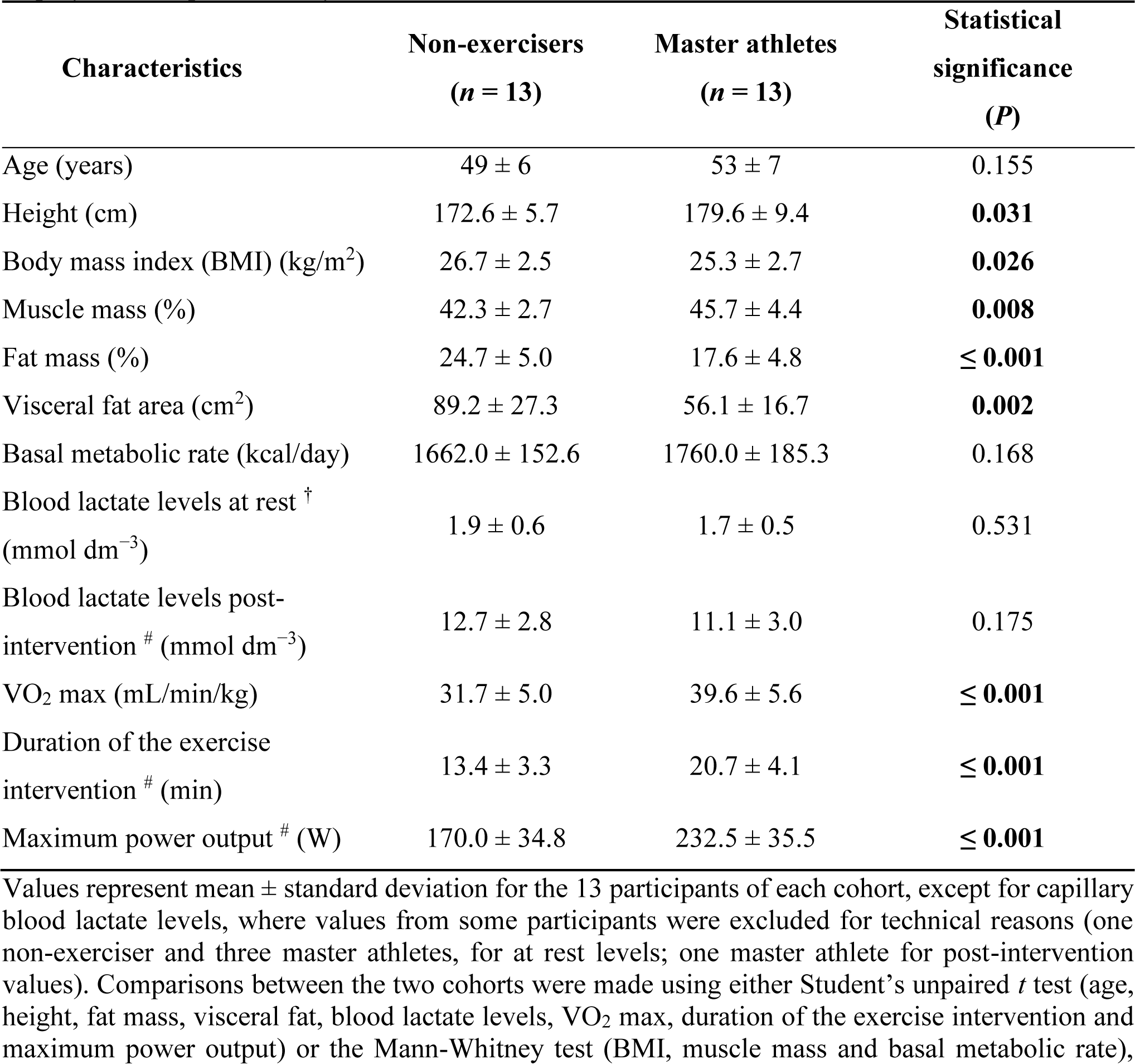
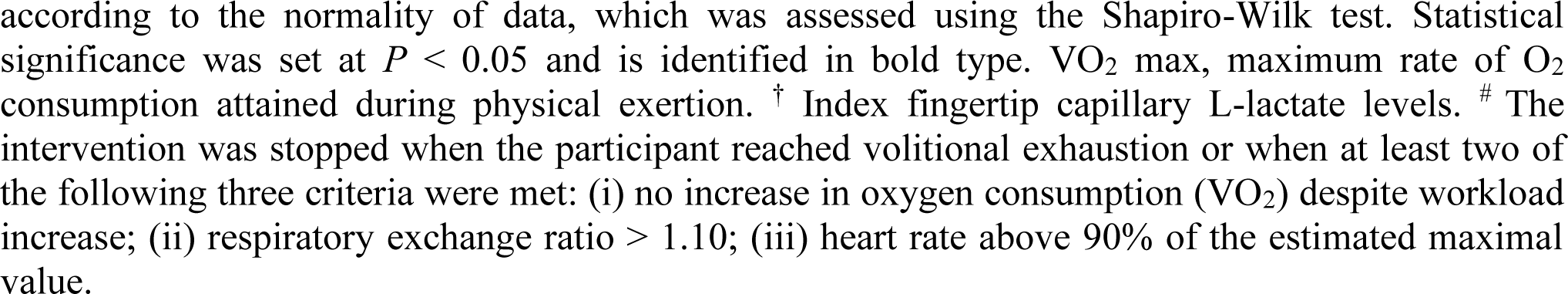
Anthropometric and physiological characteristics of the two age-matched male cohorts employed in the present study.

| Characteristics | Non-exercisers<br>(*n* = 13) | Master athletes<br>(*n* = 13) | Statistical<br>significance<br>(*P*) |
| --- | --- | --- | --- |
| Age (years) | 49 ± 6 | 53 ± 7 | 0.155 |
| Height (cm) | 172.6 ± 5.7 | 179.6 ± 9.4 | **0.031** |
| Body mass index (BMI) (kg/m<sup>2</sup>) | 26.7 ± 2.5 | 25.3 ± 2.7 | **0.026** |
| Muscle mass (%) | 42.3 ± 2.7 | 45.7 ± 4.4 | **0.008** |
| Fat mass (%) | 24.7 ± 5.0 | 17.6 ± 4.8 | **≤ 0.001** |
| Visceral fat area (cm<sup>2</sup>) | 89.2 ± 27.3 | 56.1 ± 16.7 | **0.002** |
| Basal metabolic rate (kcal/day) | 1662.0 ± 152.6 | 1760.0 ± 185.3 | 0.168 |
| Blood lactate levels at rest <sup>†</sup><br>(mmol dm<sup>-3</sup>) | 1.9 ± 0.6 | 1.7 ± 0.5 | 0.531 |
| Blood lactate levels post-intervention <sup>#</sup> (mmol dm<sup>-3</sup>) | 12.7 ± 2.8 | 11.1 ± 3.0 | 0.175 |
| VO<sub>2</sub> max (mL/min/kg) | 31.7 ± 5.0 | 39.6 ± 5.6 | **≤ 0.001** |
| Duration of the exercise<br>intervention <sup>#</sup> (min) | 13.4 ± 3.3 | 20.7 ± 4.1 | **≤ 0.001** |
| Maximum power output <sup>#</sup> (W) | 170.0 ± 34.8 | 232.5 ± 35.5 | **≤ 0.001** |

Values represent mean ± standard deviation for the 13 participants of each cohort, except for capillary blood lactate levels, where values from some participants were excluded for technical reasons (one non-exerciser and three master athletes, for at rest levels; one master athlete for post-intervention values). Comparisons between the two cohorts were made using either Student's unpaired *t* test (age, height, fat mass, visceral fat, blood lactate levels, VO<sub>2</sub> max, duration of the exercise intervention and maximum power output) or the Mann-Whitney test (BMI, muscle mass and basal metabolic rate),
according to the normality of data, which was assessed using the Shapiro-Wilk test. Statistical significance was set at $P < 0.05$ and is identified in bold type. $\text{VO}_2$ max, maximum rate of $\text{O}_2$ consumption attained during physical exertion. <sup>†</sup> Index fingertip capillary L-lactate levels. <sup>#</sup> The intervention was stopped when the participant reached volitional exhaustion or when at least two of the following three criteria were met: (i) no increase in oxygen consumption ( $\text{VO}_2$ ) despite workload increase; (ii) respiratory exchange ratio $> 1.10$ ; (iii) heart rate above 90% of the estimated maximal value.

As expected, master athletes exhibited a significantly higher cardiorespiratory fitness than non-exercisers (Rogers *et al*., 1990; Mckendry *et al*., 2018), as assessed by the maximum rate of O_2_ consumption attained during physical exertion (VO_2_ max) and by the duration of the exercise intervention, whose mean values were, respectively, *ca*. 25% (*P* ≤ 0.001) and 50% higher (*P* ≤ 0.001) than those of non-exercisers. In line with the increase in the duration of the intervention, the maximum power output increased by *ca*. 35% (*P* ≤ 0.001). Basal metabolic rate and blood lactate levels (at rest and after the exercise intervention), on the other hand, were not significantly different between the two cohorts. However, the increase in blood lactate levels produced by the exercise intervention differed substantial amongst the different participants of each cohort, with individual increases ranging between 5.2-fold and 9.5-fold, amongst non-exercisers, and between 2.5-fold and 12.1-fold amongst master athletes (data not shown). These differences did not correlate with either the duration of the exercise intervention or the maximal power output or VO_2_ max (Pearson’s determination coefficients for these hypothetical correlations, *R*^2^, ranged between 0.004 and 0.126, and the corresponding *P* values ranged between 0.081 and 0.780).

Baseline serum levels of CRP, a biomarker of inflammation (Sproston & Ashworth, 2018), were also determined for each participant. Of the 13 non-exercisers, 12 had normal CRP levels (< 0.5 mg/dL), whereas one had a slightly increased level (0.72 mg/dL). All master athletes had normal CRP levels. CRP levels were below the limit of detection of the standard CRP test employed (0.1 mg/dL) in sera from 5 non-exercisers and in sera from 10 master athletes (data not shown). None of the small differences in the cohorts’ serum levels of glucose, lipids, and albumin were statistically significant (Table 2). Also, the effect of the exercise intervention in these levels was equivalent for all analytes tested (all increased by *ca.* 10% in both cohorts; results not shown). As such, rather than a specific response to acute exercise, the increase may reflect a transient small reduction of plasma volume due to the intervention, as previously reported for both normotensive and untreated hypertensive patients (Hansen, 1968).

**Table 2.** Baseline serum levels of glucose, cholesterol, triglycerides and albumin in the two age-matched cohorts employed in the present study.

| Analyte | Non-exercisers | Master athletes | Statistical significance |
| --- | --- | --- | --- |

|  | (n = 13) | (n = 13) | (P) |
| --- | --- | --- | --- |
| Glucose (mg/dL) | 104.7 ± 24.6 | 97.6 ± 17.0 | 0.605 |
| Cholesterol (mg/dL) | 212.6 ± 32.4 | 218.1 ± 29.6 | 0.654 |
| HDL cholesterol (mg/dL) | 49.7 ± 12.2 | 57.4 ± 7.5 | 0.065 |
| LDL cholesterol (mg/dL) | 126.3 ± 27.4 | 123.4 ± 22.3 | 0.797 |
| Triglycerides (mg/dL) | 128.6 ± 34.0 | 106.4 ± 40.5 | 0.143 |
| Albumin (g/L) | 50.2 ± 7.4 | 52.1 ± 2.0 | 0.373 |

Each serum was tested twice for the different analytes and the two results were averaged. Values represent mean ± standard deviation for the averages of the 13 participants in each cohort. Comparisons between the two cohorts were made using either unpaired Student's *t*-test (cholesterol, HDL cholesterol, LDL cholesterol, triglycerides and albumin) or the Mann-Whitney test (glucose), according to the normality of data, which was assessed using the Shapiro-Wilk test. For all analytes tested, no statistically significant differences were found between the two cohorts (statistical significance was set at $P < 0.05$ ). HDL, High-density lipoprotein; LDL, Low-density lipoprotein.

### Serum conditioning by acute exercise reduced the proliferation rate of A549 cells, whereas conditioning by lifelong exercise had the opposite effect

As can be appreciated in Figure 1, acute and lifelong exercise had opposite effects on cell proliferation: whereas serum conditioned by acute exercise decreased cell proliferation by *ca.* 10% (*P* = 0.062), for non-exercisers, and by *ca.* 15% (*P* = 0.003), for master athletes, serum conditioned by lifelong exercise stimulated it by *ca*. 10% (*P* = 0.035). The effect of acute exercise was highly consistent in the masters’ cohort, being observed in 11 out of the 13 (85%) individuals, but less consistent in the non-exercisers’ cohort, where it was only observed in 8 out of the 13 (62%) individuals. Nonetheless, the decrease in proliferation rate produced by the intervention did not differ, on average, significantly between the two cohorts (*P* = 0.406).

**Figure 1.**
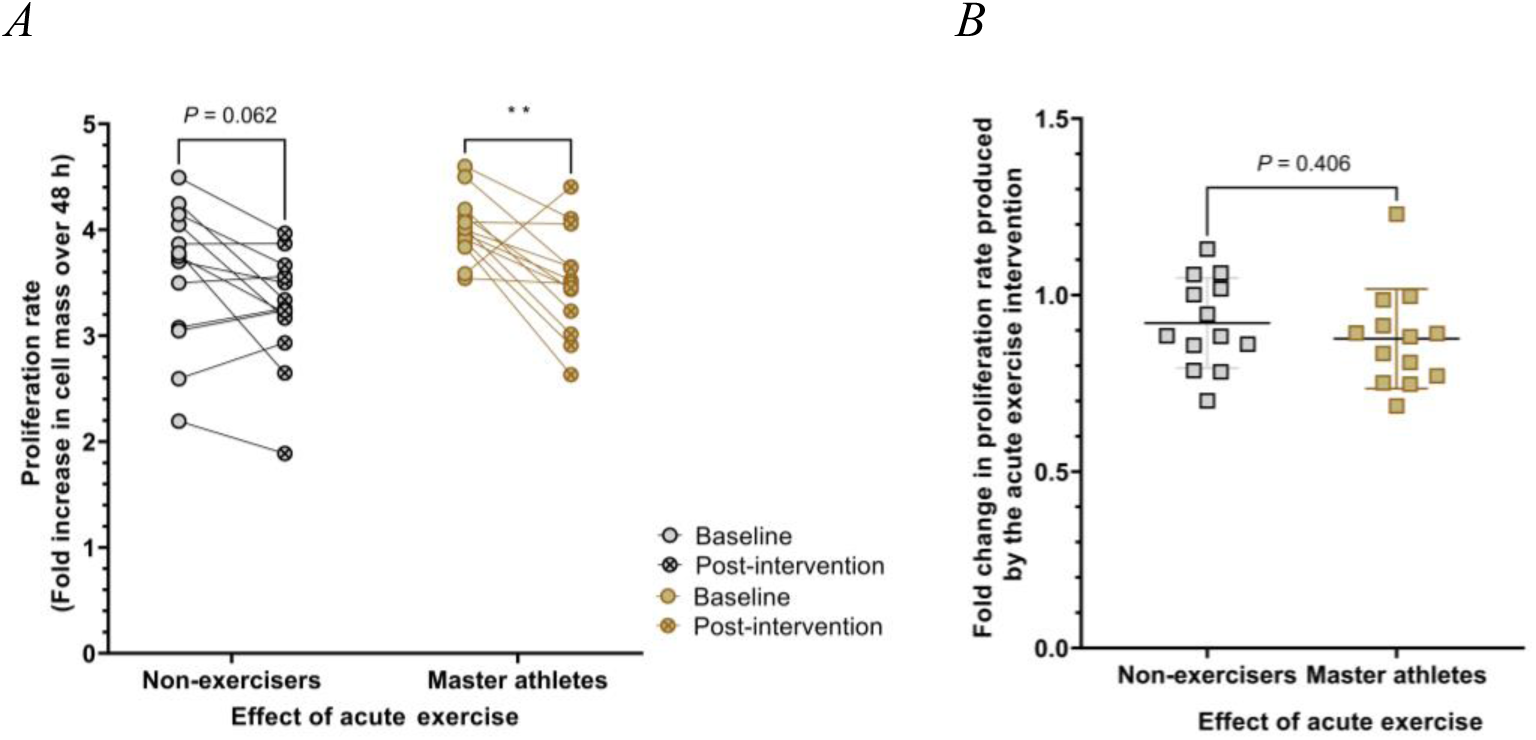
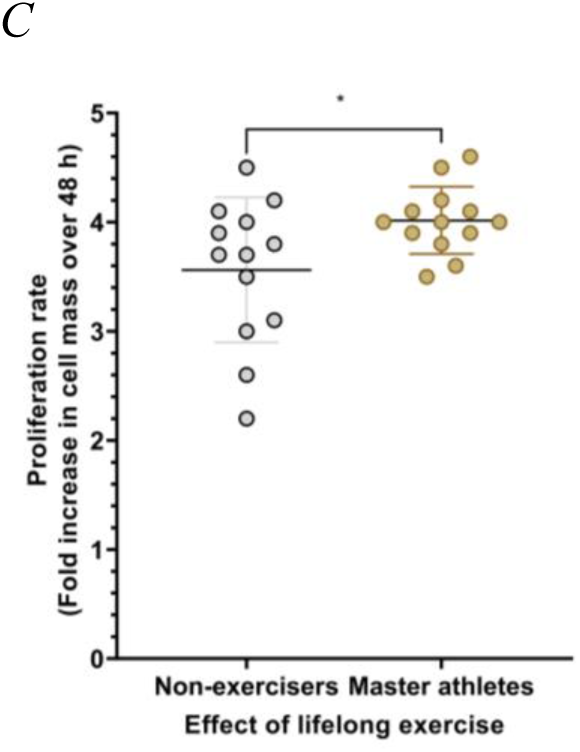
Serum conditioning by acute and lifelong exercise had opposite effects on the proliferation rate of A549 human lung cancer cells. (*A*) For most of the participants in both cohorts, conditioning by acute exercise reduced sera’s ability to stimulate cell proliferation. (*B*) On average, the impact of the acute exercise intervention on serás ability to stimulate cell proliferation did not differ significantly (ns) between the two cohorts (right panel). (*C*) On average, cells grown in medium supplemented with baseline masters’ serum exhibited higher proliferation rates than those grown in medium supplemented with baseline non-exercisers’ serum. All cultures were established and grown in medium supplemented with 10% (v/v) human serum. The number of cells in culture at 24 h and 48 h post-seeding, used to calculate proliferation rates, was estimated using the sulforhodamine B (SRB) assay. Dots and squares represent individual values for the different sera and are the means of three independent experiments. In each of these experiments, all conditions (sera and time points) were tested in three replicate cultures. Each set of connected dots represents sera collected from the same participant before (left; Baseline) and after (right; Post-intervention) the acute exercise intervention. Large horizontal bars and associated smaller horizontal (error) bars represent means ± standard deviation for the 13 participants of each cohort. The statistical significance of the effect of lifelong exercise was assessed using Student’s unpaired *t*-test. All other comparisons were made by repeated measures two-way ANOVA, assuming sphericity. Whenever the null hypothesis was rejected, Šídák’s multiple comparison test was performed. Statistical significance was set at *P* < 0.05 and statistically significant differences between the indicated groups are shown by *, for *P* ˂ 0.05, and by **, for *P* < 0.01.

### Serum conditioning by acute exercise increased the lag phase of A549 cells, whereas conditioning by lifelong exercise had the opposite effect

When assessing proliferation rates, we observed, for both cohorts, that the number of cells in culture 24 h post-seeding was, on average, ca. 20% higher in cultures stimulated by post-intervention serum than by the respective baseline serum (P ˂ 0.0001 for both cohorts; Figure 2*A,B*). Of note, this was highly consistent, being observed in 12 out of 13 (92%) non-exercisers and in all (100%) master athletes. These differences in cell number cannot be explained by different proliferation rates, as our proliferation data point in the opposite direction, i.e., cultures with the higher number of cells 24 h post-seeding had lower proliferation rates (Figure 1). Also, microscopic observation of the cultures did not reveal any differences in the number of dead or unattached cells. Therefore, cells exposed to sera conditioned by acute exercise took less time to attach to the substrate, spread and/or resume proliferation than those exposed to the corresponding baseline sera. Surprisingly, conditioning of sera by lifelong exercise had the opposite effect, resulting in ca. 10% decrease in the number of cells in culture 24 h post-seeding (P = 0.281; Figure 2*C*) suggesting and increased lag phase.

**Figure 2.**
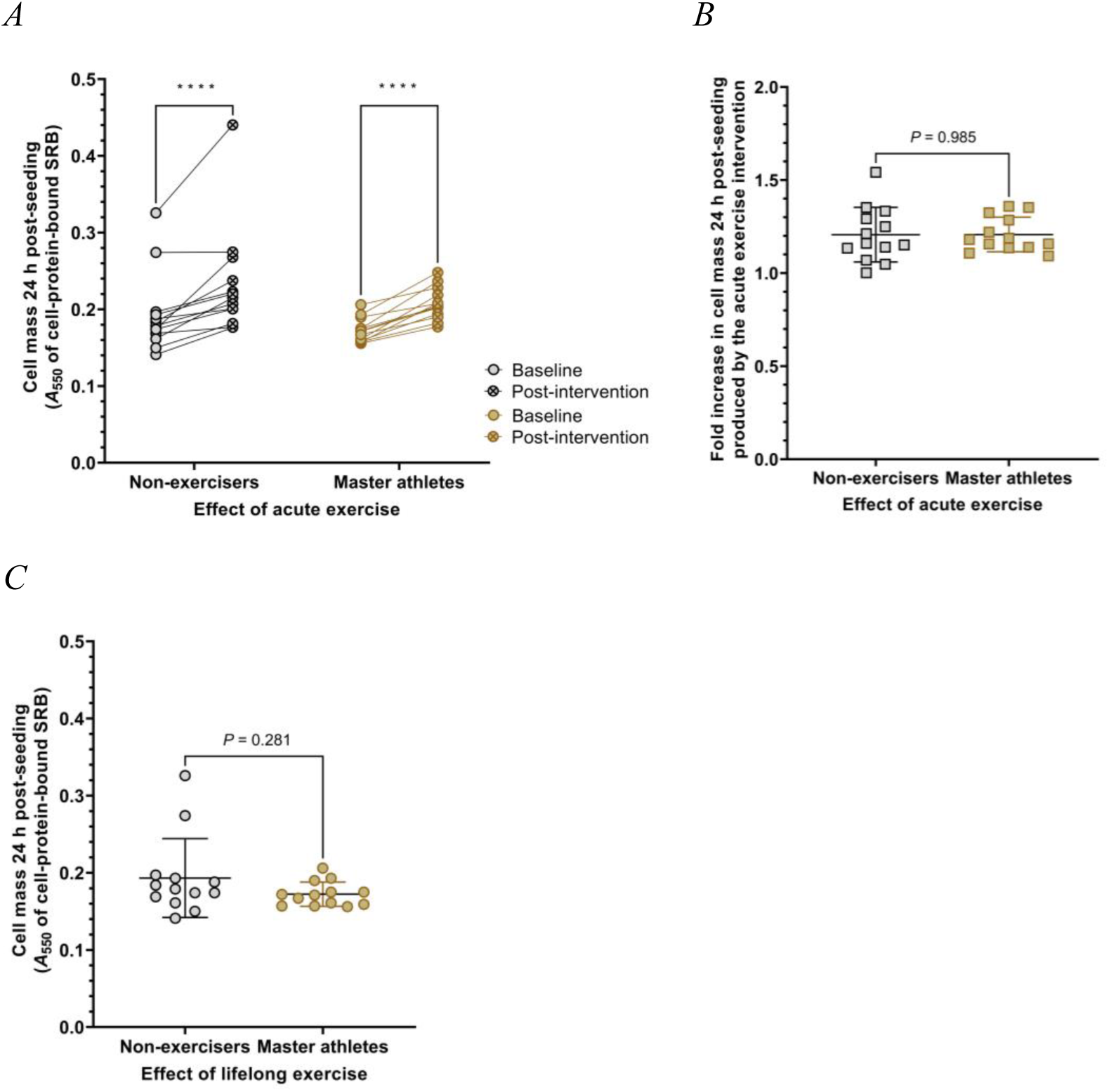
Serum conditioning by lifelong and acute exercise had opposite effects on the number of A549 human lung cancer cells in culture 24 h post-seeding. (*A*) For both cohorts, cultures grown in medium supplemented with serum conditioned by the acute exercise intervention (post-intervention serum) contained, on average, *ca*. 20% more cells 24 h post-seeding than those grown in medium supplemented with baseline serum from the same participant, suggesting that the serum changes produced by the intervention shortened the cells’ lag phase. This increase was observed in 12 out of 13 (92%) non-exercisers and in all (100%) master athletes. (*B*) No statistically significant differences (ns) were obtained between the two cohorts in terms of the impact of acute exercise serum conditioning on the number of cells in culture 24 h post-seeding. (*C*) On average, cultures grown in medium supplemented with baseline masters’ serum contained *ca.* 10% less cells 24 h post-seeding than those grown in medium supplemented with baseline non-exercisers’ serum. All cultures were established and grown in medium supplemented with 10% (v/v) human serum. The total amount of cell protein, assessed using the sulforhodamine B (SRB) assay, was used as an estimate of the number of cells in culture. Dots and squares represent individual values for the different sera and are the means of three independent experiments. In each of these experiments, all sera were tested in three replicate cultures and the values were averaged. Each set of connected dots represents baseline (left) and post-intervention (right) sera from the same participant. Large horizontal bars and associated smaller horizontal (error) bars represent means ± standard deviation for the 13 participants of each cohort. The statistical significance of the effect of lifelong exercise was assessed using the Mann-Whitney test. All other comparisons were made by repeated measures two-way ANOVA, assuming sphericity. Whenever the null hypothesis was rejected, Šídák’s multiple comparison test was performed. Statistical significance was set at *P* < 0.05 and statistically significant differences between the indicated groups are shown by ****, for *P* ˂ 0.0001. *A*_550_, Absorbance at 550 nm.

### Neither acute nor lifelong exercise serum conditioning affected the reproductive potential or the migratory capacity of A549 cells

We next investigated the impact of exercise on the ability of cells to form colonies when seeded at very low densities, *i.e.*, on their plating efficiencies, used here as a metric for reproductive potential. Our results show that neither acute nor lifelong exercise had a significative impact on this parameter (Figure 3*A*–*C*; *P* = 0.692 and *P* = 0.940, for the effect acute exercise on non-exercisers and master athletes, respectively, and *P* = 0.396 for the effect of lifelong exercise). Regarding the very large changes in colony size observed (Figure 3*D*–*F*), they are likely the result of the alterations in proliferation rates discussed above (Figure 1). It must be noted that small differences in proliferation rate, such as those observed in our study, can translate into markedly different cell numbers at the end of 9 days (the duration of the assay employed).

**Figure 3.**
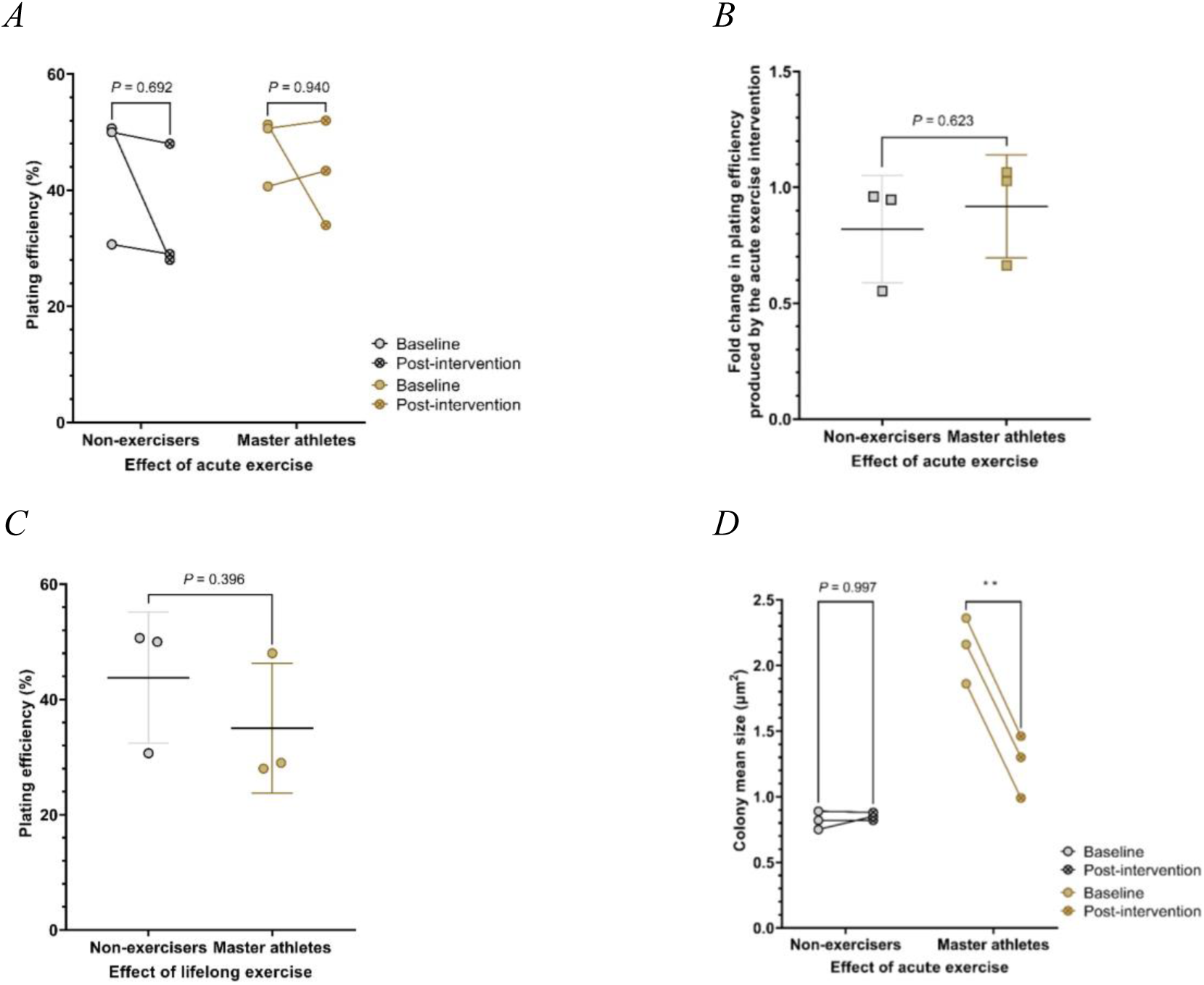
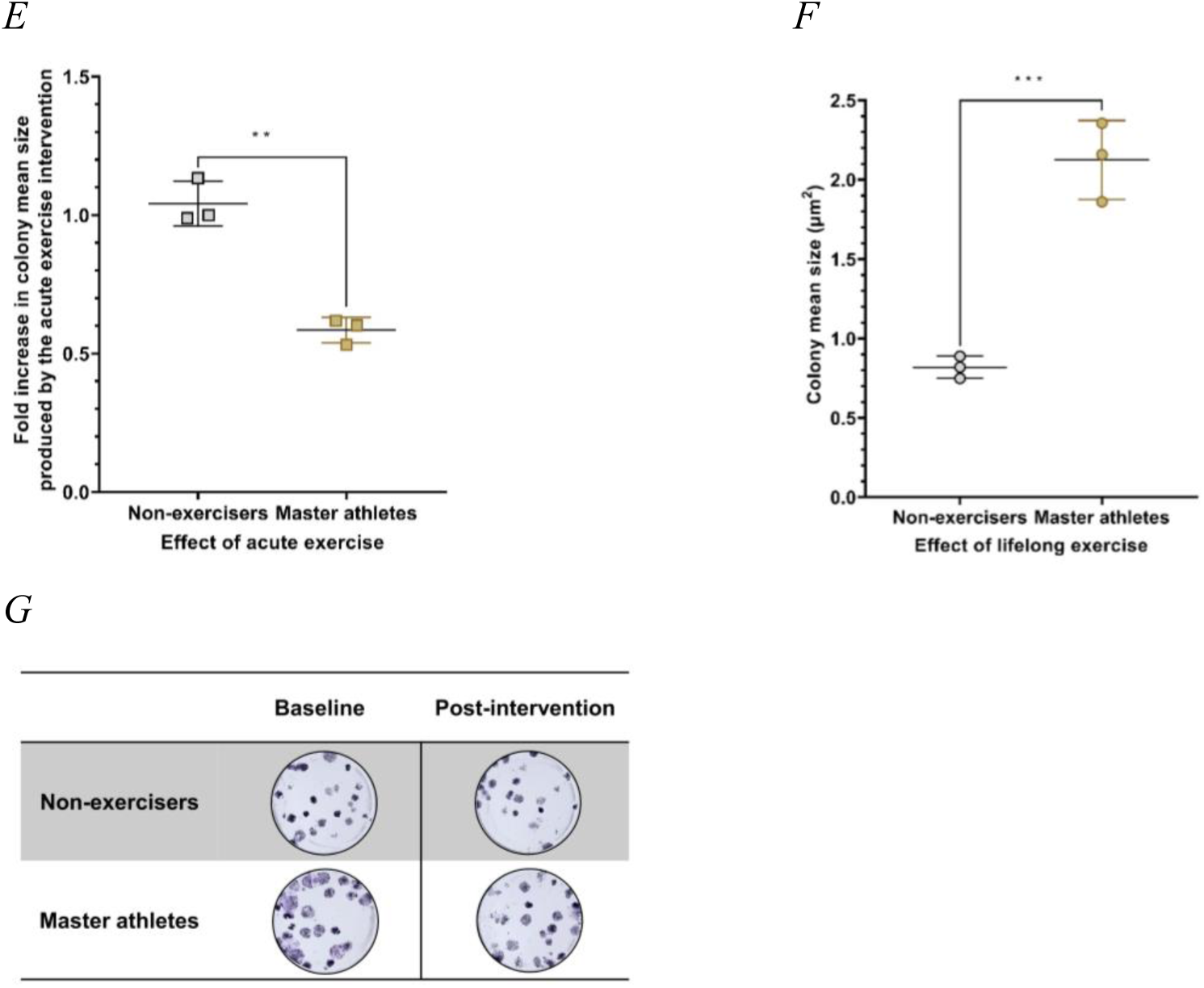
Neither acute nor lifelong exercise serum conditioning affected the reproductive potential of A549 human lung cancer cells, as assessed by their plating efficiencies. Pooled sera were used for the four conditions tested (baseline non-exercisers; baseline master athletes; post-intervention non-exercisers; post-intervention master athletes), each containing equal volumes of serum from 13 participants. (*A–C*) Plating efficiencies, expressed as percentage values, were calculated by dividing the number of colonies formed by the number of cells seeded, whereas (*D–F*) colony mean sizes were calculated dividing the area covered by the colonies (measured using ImageJ) by the total number of colonies. Each set of three dots and squares represents the values obtained in three independent experiments. In each of these experiments, all pooled sera were tested in three replicate cultures and the results averaged. Large horizontal bars and associated smaller horizontal (error) bars represent means ± standard deviation for the three independent experiments. The statistical significance of the effect of lifelong exercise was assessed using Student’s unpaired *t*-test. All other comparisons were made by repeated measures two-way ANOVA, assuming sphericity. Whenever the null hypothesis was rejected, Šídák’s multiple comparison test was performed. Statistical significance was set at *P* < 0.05 and statistically significant differences between the indicated groups are shown by **, for *P* ˂ 0.01, and ***, for *P* ˂ 0.001. ns, No statistically significant differences were obtained (i) in plating efficiency between baseline and corresponding post-intervention pooled sera in the two cohorts and between baseline sera from the two cohorts; (ii) in the mean size of the colonies between baseline and post-intervention sera in non-exercisers. (*G*) Representative photographs of the colonies that formed after 9 days of incubation when cells were plated using a single-cell suspension at a colony-forming density of 40 cells per well in 24-well plates, in 500 µL of growth medium supplemented with 10% (v/v) of pooled human serum.

The results of our cell migration assessments are summarized in Figure 4. As can be appreciated, neither acute nor lifelong exercise conditioning altered serum’s ability to promote cell migration to a statistically significant extent (Figures 4*A–C*; *P* = 0.998 and *P* = 0.555, for the effect of acute exercise on non-exercisers and master athletes, respectively; *P* = 0.448, for the effect of lifelong exercise). It must be noted, though, that cells were exposed to human sera for a relatively short period (18 h). Also, due to the limited volume of human serum available, the results are from a single experiment (in which all 52 sera were tested in duplicate).

**Figure 4.**
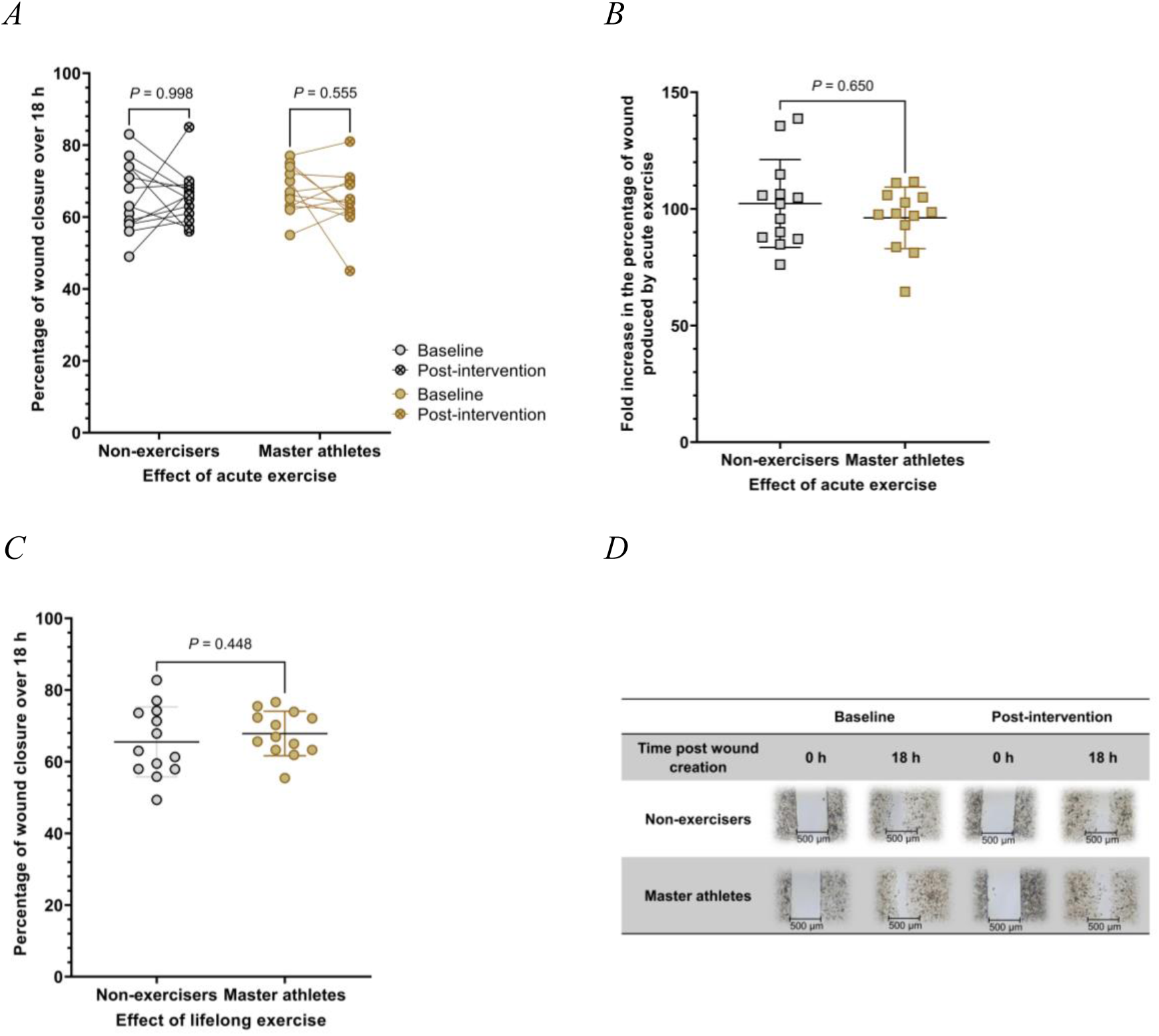
Neither acute nor lifelong exercise serum conditioning affected the *in vitro* migratory capacity of A549 human lung cancer cells. (*A–C*) To assess *in vitro* migratory capacity, two artificial 500 µm cell-free gaps were created in confluent monolayer cultures, using 3-well culture inserts that were placed onto the wells of 24-well plates, one insert per well and per serum tested. Cultures were then exposed to medium supplemented with 10% (v/v) human serum. For each gap, three different regions were imaged immediately after its creation and after an 18-hour incubation. Cell-free areas in these images were then measured using ImageJ and the percentage of gap closure over this period was calculated. For each serum, the average of the corresponding six values was then used as an estimate of cell migration potential. Dots and squares represent individual values for the different sera. Each set of connected dots represents baseline (left) and post-intervention (right) sera from the same participant. Large horizontal bars and associated smaller horizontal (error) bars represent means ± standard deviation for the 13 participants of each cohort. The statistical significance of the effect of lifelong exercise was assessed using the Mann-Whitney test. All other comparisons were made by repeated measures two-way ANOVA, assuming sphericity. Statistical significance was set at *P* < 0.05 and the null hypothesis was never rejected (ns). Results are from a single independent experiment in which each serum was tested in two artificial gaps. (*D*) Representative micrographs (40× magnification) of artificial gaps for each of the four conditions tested, taken immediately after the creation of the gap and after an 18-h incubation in growth medium supplemented with 10% (v/v) human serum. Micrographs were captured using an Olympus CKX53 inverted optical microscope equipped with a camera and the EPview™ software (V2.9.6_20201224; Hachioji, Tokyo, Japan).

### Lifelong exercise increased the sensitivity of A549 cells to cisplatin

As can be appreciated in Figure 5, sensitivity to cisplatin was much higher (as assessed by the lower CC_50_ value) when cells were incubated in the presence of baseline serum from master athletes, than baseline serum from non-exercisers (*P* = 0.04). On the contrary, acute exercise had no significant impact on this sensitivity (*P* = 0.999 and *P* = 0.712, for non-exercisers and master athletes, respectively).

**Figure 5.**
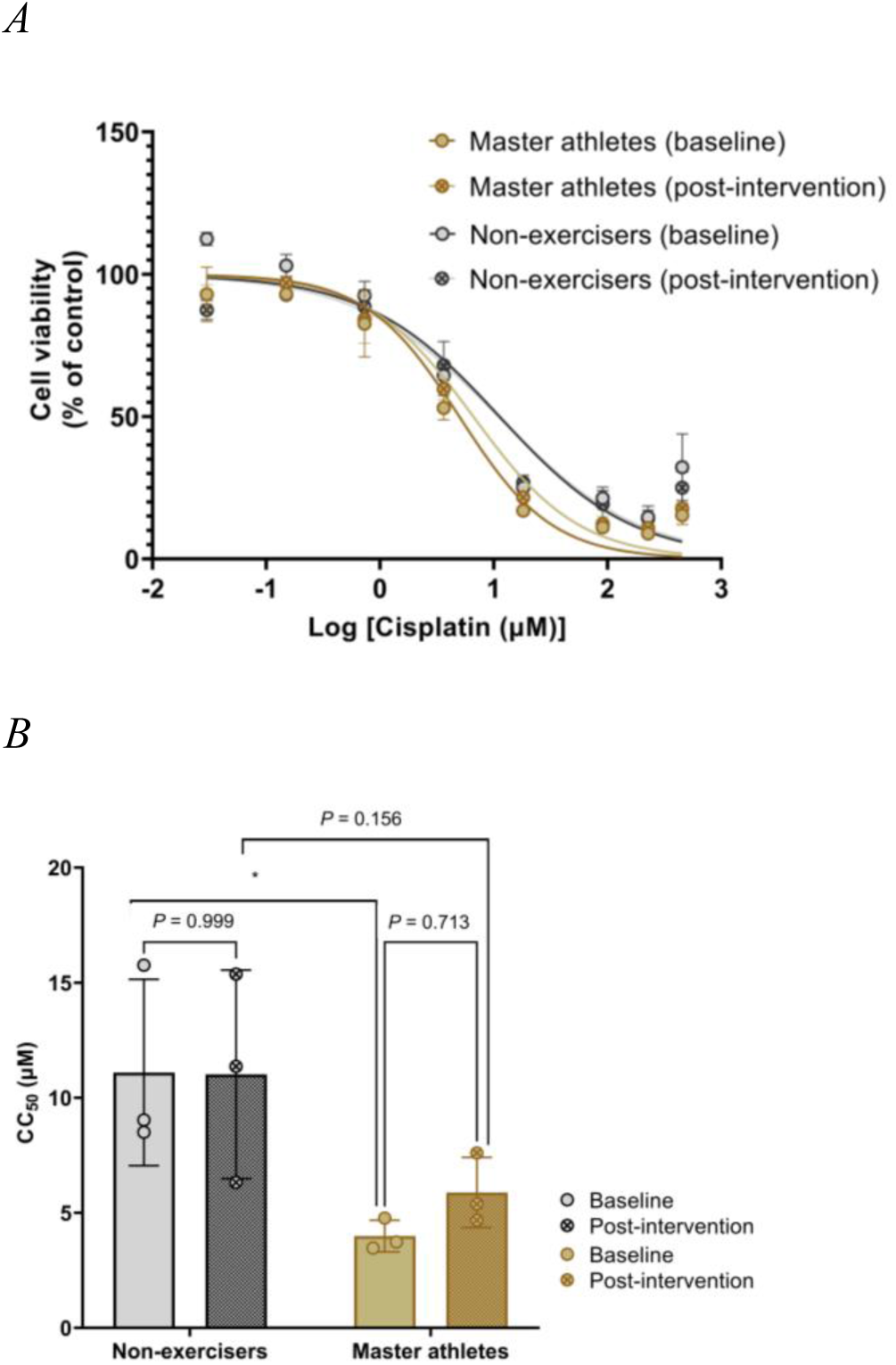
Lifelong exercise more than doubled the sensitivity of A549 human lung cancer cells to cisplatin. (*A*) Concentration-response curves depicting the cytotoxicity of cisplatin against cells stimulated with pooled sera (10% (v/v)) from non-exercisers and master athletes, obtained at rest (Baseline) and after an acute exercise intervention (Post-intervention). (*B*) CC_50_ values for cisplatin for cultures incubated with the four different pooled sera. Cell viability was estimated at 72 h post-cisplatin addition using the sulforhodamine B (SRB) assay. For each pooled serum, the impact of the different cisplatin concentrations (0–455 μM) on cell viability was tested in three independent experiments, each with three replicate cultures per concentration, and expressed as percentage of the control value (0 µM cisplatin). CC_50_ values were calculated from concentration-response curves generated using a nonlinear regression model in GraphPad Prism. Dots/diamonds and corresponding error bars represent, respectively, means and standard deviation for the three independent experiments. Comparisons between the indicated groups were made by both (*A*) the sum-of-squares using the extra sum-of-squares F test; (*B*) Student’s unpaired t-test (effect of lifelong exercise) and repeated measures two-way ANOVA, assuming sphericity (effect of acute exercise). Whenever the null hypothesis was rejected, Šídák’s multiple comparison test was performed. Statistical significance was set at P < 0.05 and statistically significant differences between the indicated groups are shown by *, for P ˂ 0.05, and by **, for P < 0.01. CC_50_, Half-maximal cytotoxic concentration.

### Both acute and lifelong exercise modulated serum cytokine levels

Acute exercise increased the levels of most cytokines (*ca.* 50 in both cohorts) and these increases tended to be substantial (up to 7-fold). Unlike in non-exercisers’ serum, where there was a very large increase in IL-1β, IL-2 and IL-15, among others, and a very large decrease in IL-6, among others, following acute exercise, no effect was observed in the levels of these cytokines in master athletes’ serum. Interestingly, for some cytokines, such as GRO (α/β/γ), IL-8, IL-10 and MIP-1β, acute exercise had opposite effects in the two cohorts, reducing their levels in non-exercisers, but augmenting them in master athletes (Figure 6*A*).

**Figure 6.**
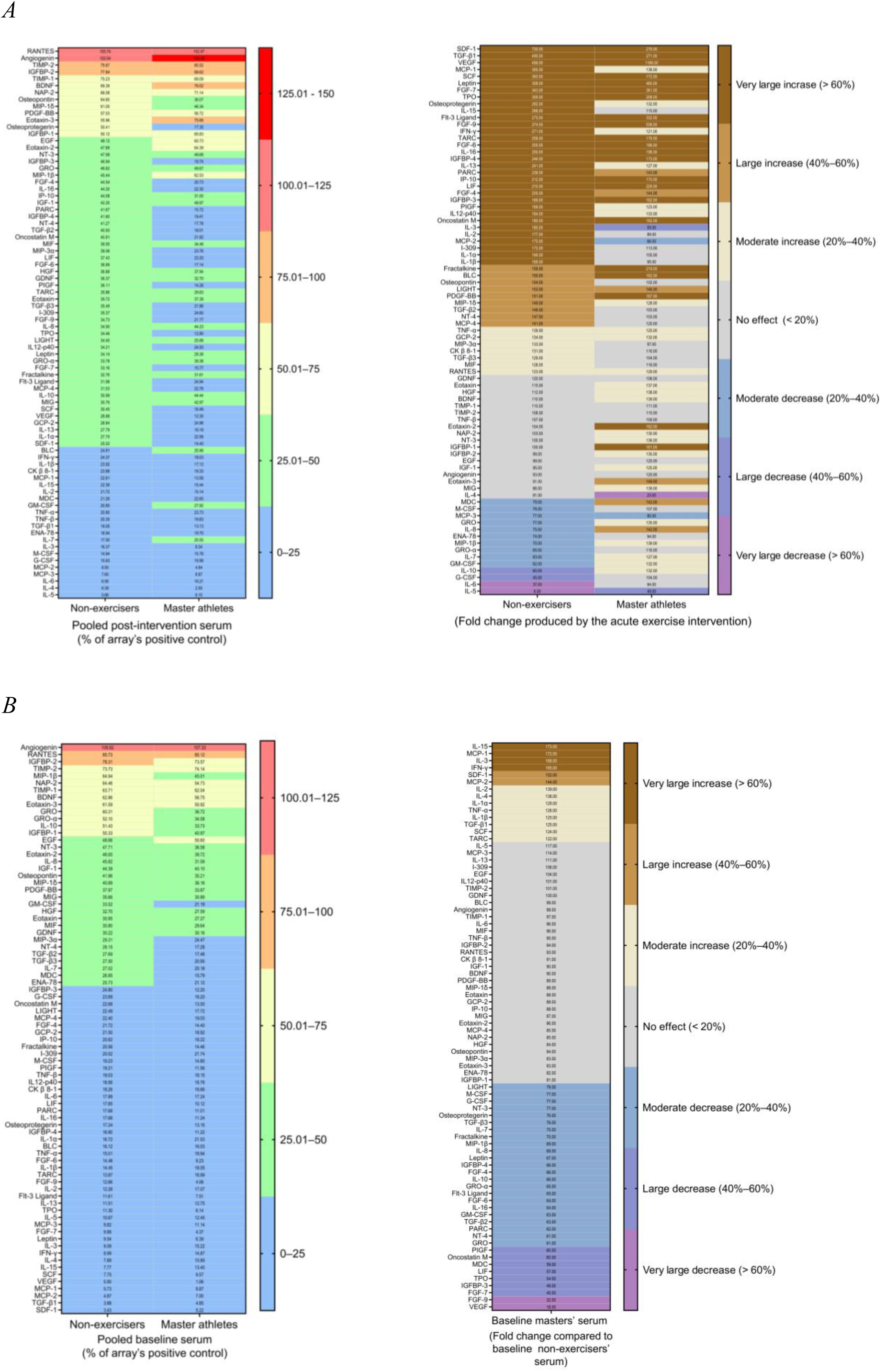
The effects of exercise on serum cytokine levels depended on cytokine, duration of the exercise (acute versus lifelong) and previous levels of physical activity (non-exercisers versus master athletes). The semi-quantitative assessment of the 80 cytokines was carried out in pooled sera, each containing equal volumes of serum from 13 participants, using an antibody-based microarray. For each antigen-specific antibody spot, chemiluminescence signal intensity (average pixel/area) was determined using ImageJ. Two independent determinations were carried out and the results were averaged (*A,B*; Left) Cytokine levels are presented in decreasing order of signal intensity in the pooled non-exercisers’ serum. (*A,B*; Right) Cytokines are presented in order of fold change in signal intensity produced by the acute exercise intervention in the non-exercisers’ serum.

Regarding lifelong exercise, it had no effect on *ca.* 40% of the cytokines tested and when it did have an effect it was mostly a decrease: 32 cytokines saw their signal intensity decrease. The 14 cytokines increased, most notably MCP2, SDF1, and TGF-β1 (Figure 6*B*), displayed very weak signals near the limit of detection in both cohorts and their increase may thus not represent true physiological changes. On the contrary, many of the cytokines whose signals were significantly reduced in the serum of master athletes compared to the serum of non-exercisers, namely GM-CSF, GRO (α/β/γ), GRO-α, IL-8 (CXCL8), IL-10, MIP-1β (CCL4), NT-3 and osteopontin, were significantly expressed in both cohorts.

## Discussion

Over 30% of the worldwide population does not attain the levels of physical activity for health and well-being recommended by the WHO (Bull *et al*., 2020), (Elgaddal *et al*., 2022). Establishing a causal relationship between increased levels of physical activity and lower cancer risk and mortality would constitute a strong inducement for inactive people to significantly reduce the amount of time spent in sedentary occupations. Towards this goal, several research groups exposed cultures of human cancer cell lines to human serum obtained before and after an acute bout of exercise to determine whether the transient systemic changes induced directly impacted their behavior (Rundqvist *et al*., 2013; Dethlefsen *et al*., 2016; Dethlefsen *et al*., 2017; Kurgan *et al*., 2017; De Santi *et al*., 2019; Devin *et al*., 2019; Baldelli *et al*., 2020; Hwang *et al*., 2020; Orange *et al*., 2022; Kim *et al*., 2023). These studies focused mostly on cell proliferation, which was often assessed by the number of cells in culture after a specified period of incubation. Other parameters were also investigated, but not as consistently, namely apoptosis levels (Leung *et al*., 2004; Rundqvist *et al*., 2013; Devin *et al*., 2019) and the cells’ ability to form colonies when seeded at very low densities (Kurgan *et al*., 2017) or under conditions that prevent cell attachment (De Santi *et al*., 2019; Baldelli *et al*., 2020). Despite the scarcity of data and significant inter-study heterogeneity, meta-analysis of the results indicates that serum conditioning by acute exercise significantly reduced cancer cell proliferation, with a large effect size that increased with the intensity of the exercise (Soares *et al*., 2021). Of note, for all cell lines/cohorts reported (Rundqvist *et al*., 2013; Dethlefsen *et al*., 2016; Dethlefsen *et al*., 2017; Kurgan *et al*., 2017; Devin *et al*., 2019; Baldelli *et al*., 2020; Hwang *et al*., 2020; Orange *et al*., 2022; Kim *et al*., 2023), only in one single case (PC3 cells and a young age cohort) did acute exercise failed to decrease cell proliferation (Hwang *et al*., 2020). Significantly, in a study that employed both normal (MRC5) and cancer (A549) lung cell lines, the proliferation of normal cells was not affected, while that of cancer cells was (Kurgan *et al*., 2017).The effects of chronic exercise on the properties of cultured human cancer cells, on the other hand, have not yet been the subject of meta-analysis, due to the reduced number of studies (Barnard *et al*., 2003; Leung *et al*., 2004; Dethlefsen *et al*., 2016; Baldelli *et al*., 2020; Hwang *et al*., 2020; Kim *et al*., 2022). Also, the duration of the exercise intervention in most of these studies was relatively short (1 to 6 months) (Dethlefsen *et al*., 2016; Baldelli *et al*., 2020; Hwang *et al*., 2020; Kim *et al*., 2022), with only two studies exploring the effects of exercise training over 10 or more years (Barnard *et al*., 2003; Leung *et al*., 2004). Notwithstanding the scarcity of data, the results from these studies suggest that the impact of chronic exercise on the ability of sera to stimulate cancer cell proliferation differs from that of sera conditioned by acute exercise (see below).

The aim of the present investigation was three-fold. First, to strengthen the evidence regarding the impact of exercise conditioning on the proliferation of cultured human cancer cells. Second, to expand this line of investigation to cell migration, a cell property also frequently used as a marker of transformation degree and, ultimately, tumorigenic potential, which was not previously investigated in this type of study. Compromising or losing any of these markers might ultimately halt or reverse cancer progression. Last, but not least, to investigate, for the first time, whether the decreased cancer mortality associated with exercise might be partially explained by an increased sensitivity of cancer cells to chemotherapy agents. Our study employed two cohorts of age-matched individuals that differed significantly in terms of levels of physical activity, specifically individuals who did not meet the WHO guidelines on physical activity and master athletes (as a model of lifelong exercise). The simultaneous use of two distinct cohorts allowed to gain insight into the dependency of the cellular effects induced by acute exercise on cohort characteristics.

All cell studies were carried out on cultures of A549 cells, which were used as an *in vitro* model of human lung cancer, for which epidemiological evidence linking physical activity with reduced cancer risk is strong (Moore *et al*., 2016). Cellular studies are prone to a high degree of variability, due namely to intra-cell-line heterogeneity. To increase the robustness of our data, multiple biologically independent experiments were carried out, and each serum was tested on at least two technical replicates on each of these experiments.

For our analysis of the impact of exercise on proliferation rate, we employed the SRB assay (Vichai & Kirtikara, 2006). This assay quantifies total cellular protein, from which cell numbers and, ultimately, cell proliferation can be estimated. The number of cells in culture at any given time after seeding depends on parameters other than proliferation rates, namely levels of cell death and the time cells take to adhere to the substrate, spread and resume growth and division (*i.e.*, the duration of the cells’ lag phase). To omit the lag phase from our estimation of proliferation rates, cell numbers were assessed at two time points (24 h and 72 h post-seeding), rather than just at the end of the experiment, as was frequently the case in similar studies. Proliferation was then estimated as the fold increase in the number of cells over the 48-h period. Also, instead of establishing cultures in growth medium supplemented with FCS and only exposing them to human serum 24 h post-seeding, as was also often the case, our cultures were established in growth medium supplemented with human serum, *i.e.*, they were exposed to human serum throughout the experiment. This protocol not only yielded more reliable proliferation data, but also enabled us to gain insight into the influence of exercise conditioning on the lag phase.

Our results support previous findings that acute exercise decreases cancer cell proliferation (Figure 1*A,B*). Lifelong exercise, on the other hand, increased cell proliferation (Figure 1*C*). This contrasts with the reports that 10+ years of training decreased the proliferation of LNCaP prostate cancer cells, as determined by end-point analysis, and did not alter the proliferation of a LNCaP-derived cell line with nonfunctional p53 (LN-56 cells), suggesting that the effects on cell proliferation depend on the cells’ characteristics, namely their p53 status (Barnard *et al*., 2003; Leung *et al*., 2004). It also contrast with the results of three of the studies involving short-term exercise programs (1–6 months), where no effect on proliferation was observed (Dethlefsen *et al*., 2016; Devin *et al*., 2019; Baldelli *et al*., 2020; Kim *et al*., 2022). Significantly, all these three studies also assessed acute exercise, which decreased cell proliferation. This contrasting pattern between the effects of acute and chronic exercise led the authors to hypothesize that the putative beneficial effects of long-term exercise training on cancer incidence, recurrence and survival result from the cumulative effects of transient serum changes in response to each repeated exercise bout, rather than from permanent serum alterations produced by exercise training (Dethlefsen *et al*., 2016; Devin *et al*., 2019; Baldelli *et al*., 2020). However, it must be stressed that the exercise programs lasted 9 weeks at the most, which might not have been sufficient to produce permanent serum changes to factors affecting cell proliferation.

A contrasting pattern was also found in terms of the impact of the two different types of exercise on the lag phase, with acute exercise decreasing it (Figure 2*A,B*), and lifelong exercise extending it (Figure 2*C*). As the present study was the first to investigate the effects of exercise on this property, it is not possible to tell whether this effect is specific to the A549 cell line and/or the cohorts used, or a more general one. It is known that interactions between cells and the extracellular matrix (ECM) modulate cell survival and proliferation (Frantz *et al*., 2010). However, considering the limited evidence available, it would not be advisable to speculate on possible consequences of such effect on the cells’ tumorigenic potential. In addition, it must be acknowledged that, when it comes to cancer, adhesion to the ECM can be seen as a double-edged sword. In fact, in order to metastasize, cancer cells must first detach from neighboring cells and the ECM and then resist anoikis during their migration to other parts of the body. But they then need to reattach and resume proliferation, to form tumors in new locations (Welch & Hurst, 2019; Shaw *et al*., 2025). In future studies, it would be important to investigate the ability of exercise to alter the expression of adhesion molecules, such as cadherins and integrins, and the physicochemical properties of the ECM.

Neither acute (Figure 3*A,B*) nor lifelong exercise (Figure 3*C*) produced significant changes in plating efficiency, indicating that the reproductive potential was unchanged. This finding contrasts with a previously reported significant decrease in the number of colonies formed by A549 and two other cell lines when sera were conditioned by acute exercise (Kurgan *et al*., 2017). Once again, these distinct outcomes might be due to differences in study design, namely in the criteria used for counting colonies.

Together with cell–cell adhesion and cell adhesion to the substrate, the migratory capacity is often used as a metric for metastatic potential (Mehanna *et al*., 2025). In our study, neither acute (Figure 4*A,B*) nor lifelong exercise (Figure 4*C*) altered the migratory capacity of A549 cells. However, cells were exposed to human serum for a relatively short period, and it cannot be excluded that a longer exposure could have produced a different outcome.

A most striking result of our study was the strong impact that lifelong exercise had on the sensitivity to cisplatin, which more than doubled (Figure 5). Thus, one of the mechanisms by which exercise might decrease cancer mortality, besides improved immunosurveillance (Bigley *et al*., 2014; Moro-García *et al*., 2014), is by increasing the cytotoxicity of chemotherapy agents against cancer cells, thus improving therapeutic efficacy for patients. This is the first report of this type of effect and, due to its significance, this approach should undoubtedly be further pursued.

Aiming to shed light on the molecular mechanisms behind the distinct impacts of acute and lifelong serum conditioning on the behavior of cancer cells, we interrogated cytokine levels in sera of non-exercisers and master athletes at baseline and after acute exercise (Figure 6). It is well known that skeletal muscle acts as an endocrine organ that produces and secretes hundreds of cytokines and other signaling peptides, namely in response to exercise. Collectively known as myokines, these molecules mediate communication within skeletal muscle and between skeletal muscle and other organs, playing a wide array of functions (Severinsen & Pedersen, 2020). Myokines such as decorin, irisin, oncostatin M and SPARC have been implicated in the preventive and therapeutic effects of exercise and can directly affect cancer cell behavior by several means, namely by inhibiting proliferation, promoting apoptosis and inhibiting epithelial to mesenchymal cells transition, ultimately limiting invasion and metastasis. Myokines can also reduce cancer risk by indirect means, namely by inhibiting fat accumulation (*e.g.*, IL-6, IL-15, irisin and SPARC), reversing insulin resistance, reducing chronic inflammation and through modulation of the immune system (Kim *et al*., 2021; Huang *et al*., 2022).

Direct comparisons of the present work with other studies or between our two cohorts are not straightforward, as it has been found that exercise-mediated cytokine secretion depends on the characteristics of the exercise, such as type, intensity and duration. For instance, in the case of some cytokines, a significant response was only observed with more than 30 minutes of sustained moderate-to-high-intensity exercise involving major muscle groups (Piccirillo, 2019; Kim *et al*., 2021). Also, cytokine serum levels are notoriously small (in the picomolar or femtomolar range) (Anderson & Anderson, 2002). Accordingly, chemiluminescent signals for some cytokines were very weak. Changes in signal intensity produced by exercise below 20% were thus regarded as not significant. Also, increases/decreases in signal intensity above 20% must be interpreted with care in the case of cytokines whose signals were very weak in the two conditions under comparison, as small changes in signal intensity translate into marked fold changes, which do not necessarily have biological relevance. Nonetheless, some comments may be tentatively made. Namely that the effects on the serum levels of many cytokines were dependent on the type of exercise (acute versus lifelong) and cohort, as was the case with the observed effects on cell growth characteristics. Also, most of the cytokines that were affected by acute exercise had their levels increased, whereas the opposite was true for lifelong exercise. Of note, despite the changes it induced, lifelong exercise did not substantially alter the pattern of signal intensity: cytokines present at high levels in non-exercisers were also highly expressed in master athletes, and the same was true for minimally expressed cytokines. Globally speaking, the increases produced by acute exercise were more pronounced in non-exercisers than in the master athletes. For some cytokines, such as IL-8, GRO (α/β/γ), and MIP-1β, chemokines responsible for recruiting neutrophils and monocytes to sites of inflammation (Fujiwara *et al*., 2002; Parekh *et al*., 2019), and IL-10, responsible for the anti-inflammatory effect of exercise (Gleeson *et al*., 2011), acute exercise had opposite effects in the two cohorts, reducing their serum levels in non-exercisers, but augmenting them in master athletes. Of note, serum levels of several cytokines linked to the senescence-associated secretory phenotype (Li *et al*., 2023) were lower in master athletes than in non-exercisers, namely those of eotaxin-3 (CCL26), GM-CSF, GRO, IL-8, MIP-1β (CCL4) and MIP-1δ. Indeed, upon acquisition of this phenotype, fibroblasts become proinflammatory cells with the ability to promote tumor progression (Coppé *et al*., 2010).

## Conclusions

Our study is the first to address the impact of exercise on the susceptibility of cultured cancer cells to a chemotherapy drug (cisplatin), as well as on a series of properties that characterize the growth pattern of cultured human cancer cells, namely migratory capacity and cell adhesion to the substrate, resumption of cell growth and division after detachment. Other strengths of our study include the parallel investigation of the effects of both transient and permanent serum responses to, respectively, acute and lifelong exercise. Of note, we are the first to examine master athletes in this type of study and to assess the effects of acute exercise on two cohorts differing significantly in their baseline exercise levels in parallel.

In summary, we found, for both cohorts, that acute exercise decreased the proliferation rate of A549 cells, as previously reported by several groups for this and other cell lines. Interestingly, while acute exercise decreased the proliferation rate, it shortened the time cells took to adhere to the substrate, spread and resume proliferation. Lifelong exercise had the opposite effects: it increased the proliferation rate yet increased the lag phase. Regarding plating efficiency and migratory capacity, no effect could be detected. A different outcome for migratory capacity might have been observed if the duration of the exposure to human serum had been increased. Strikingly, lifelong exercise more than doubled the sensitivity of cancer cells to cisplatin, which may explain, together with other mechanisms such as enhanced immunosurveillance, the lower cancer mortality rates found among those who exercise regularly. Moreover, serum cytokine patterns induced by lifelong exercise and acute exercise contrasted sharply, potentially contributing to the different impacts that these two types of exercise had on cancer cell properties.

## Author Contributions

conceptualization, C.M.S., L.M.R.F., A.M.T. and A.M.U.; formal analysis, C.M.S., J.P.M. and A.M.U.; investigation, C.M.S., J.P.M. and A.M.U.; resources, L.R., A.P., A.M.T. and A.M.U.; writing—original draft preparation, A.M.U.; writing—review and editing, C.M.S., L.M.R.F., A.M.T. and A.M.U.; visualization, C.M.S.; supervision, A.M.T. and A.M.U.; funding acquisition, A.M.U. All authors have read and agreed to the published version of the manuscript.

## Acknowledgments

The authors wish to thank all study participants, who generously volunteered their time and efforts in participating in this study, and Dr Paulo J. Oliveira (Mitochondrial Toxicology and Experimental Therapeutics Laboratory, Center for Neuroscience and Cell Biology, University of Coimbra, Portugal), for the generous gift of the A549 cell line.

## Institutional Review Board Statement

The study was conducted in accordance with the Declaration of Helsinki and approved by the Ethics Committee of the Faculty of Sports Sciences and Physical Education of the University of Coimbra (reference CE/FCDEF-UC/00062013).

